# Error processing in the adolescent brain: Age-related differences in electrophysiology, behavioral adaptation, and brain morphology

**DOI:** 10.1101/487959

**Authors:** Knut Overbye, Kristine B. Walhovd, Tomáš Paus, Anders M. Fjell, Rene J. Huster, Christian K. Tamnes

## Abstract

Detecting errors and adjusting behaviour appropriately are fundamental cognitive abilities that are known to improve through adolescence. The cognitive and neural processes underlying this development, however, are still poorly understood. To address this knowledge gap, we performed a thorough investigation of error processing in a Flanker task in a cross-sectional sample of participants 8 to 19 years of age (n = 98). We examined age-related differences in event-related potentials known to be associated with error processing, namely the error-related negativity (ERN) and the error positivity (Pe), as well as their relationships with task performance, post-error adjustments and regional cingulate cortex thickness and surface area. We found that ERN amplitude increased with age, while Pe amplitude remained constant. A more negative ERN was associated with higher task accuracy and faster reaction times, while a more positive Pe was associated with higher accuracy, independently of age. When estimating post-error adjustments from trials following both incongruent and congruent trials, post-error slowing and post-error improvement in accuracy both increased with age, but this was only found for post-error slowing when analysing trials following incongruent trials. There were no age-independent associations between either ERN or Pe amplitude and cingulate cortex thickness or area measures.

## 1. Introduction

The abilities to detect and react to errors and to adjust ensuing behavior accordingly are fundamental for goal directed actions. Previous studies indicate that this set of abilities shows protracted development and is supported by neural structures that continue developing throughout adolescence (Tamnes et al., 2013). The error-related negativity (ERN) and error positivity (Pe) are two electrophysiological markers of central aspects of error processing. In the present study, we examined developmental age-related differences in the ERN and the Pe, and their associations with task performance, post-error adjustments and regional thickness and surface area of the cingulate cortex, a possible generator of these electrophysiological indices (Agam et al., 2011). The aim of this study was to examine how the error processing system develops from childhood to adulthood.

The ERN is a negative deflection that can be detected on the scalp usually after someone makes an incorrect response on a task (Falkenstein et al., 1991; Gehring et al., 1993; Hajcak et al., 2003). It is maximal at fronto-central recording sites, begins around the time of response, and peaks approximately 50-100 ms after an erroneous response is initiated. It is independent of the modality of stimulus and response (Ullsperger et al., 2014), and has been suggested to reflect the detection and processing of cognitive conflict, including conflict resulting from errors (Botvinick et al., 2001; Botvinick et al., 2004; Carter & Van Veen, 2007; Yeung et al., 2004), or an evaluative function signifying “worse than expected events” (Holroyd & Coles, 2002; Holroyd et al., 2005). The ERN is implicated in threat-related evaluation of errors, and is greater when errors are evaluated as more threatening (Weinberg et al., 2012), and a high ERN amplitude has been associated with higher levels of dispositional anxiety (Cavanagh & Shackman, 2015; Weinberg et al., 2016). In electrophysiological terms, it likely reflects either an increased power in the theta band and/or phase-locking of theta oscillations. The ERN is part of a family of fronto-centrally negative ERPs—including the N200 and feedback-related negativity—which are all elicited by events that trigger the need for increased cognitive control (Cavanagh & Frank, 2014; Gruendler et al., 2011; Van Noordt et al., 2016). The degree to which these ERPs functionally differ, or whether they all reflect the same latent process of performance monitoring, is a matter of debate (Gruendler et al., 2011; Van Noordt et al., 2016). The Pe is a slower, centro-parietal positive deflection that peaks between 200 and 500 ms after an incorrect response (Falkenstein et al., 2000; Gehring et al., 2012; Ladouceur et al., 2007). The functional significance of the Pe is disputed (Ferdinand & Kray, 2014). It has been suggested, for instance, that it reflects conscious awareness of making an error (Nieuwenhuis et al., 2001; Ridderinkhof et al., 2009), or the motivational or emotional significance of the error (Falkenstein et al., 2000; Overbeek et al., 2005; Ridderinkhof et al., 2009). The Pe has many similarities with the P300, which is also a centro-parietal positive deflection, related to the rapid allocation of conscious attention in tasks requiring stimulus discrimination (Polich, 2007). Davies et al. (2001) suggested that the Pe and P300 are identical, with the Pe being a P300 response to the internal detection of errors.

The body of evidence on ERN development indicates that its amplitude changes through childhood and adolescence, while other characteristics, such as its latency and scalp distribution, remains relatively constant (Grammer et al., 2014; Tamnes et al., 2013). Several cross-sectional studies have found the peak ERN amplitude to be more negative with higher age in children and/or adolescents (Davies et al., 2004a; Davies et al., 2004b; Ladouceur et al., 2004; Santesso & Segalowitz, 2008; Santesso et al., 2006). Recently, a longitudinal study also found the ERN to become increasingly negative from late childhood through adolescence (Taylor et al., 2018). Other studies, however, have found no relationship between ERN amplitude and age during this developmental period (Eppinger et al., 2009; Richardson et al., 2011). As for the Pe, a smaller number of cross-sectional studies indicate that its amplitude increases through early childhood (Grammer et al., 2014), but that it is stable from late childhood to adulthood (Davies et al., 2004b; Ladouceur et al., 2004; Wiersema et al., 2007).

The ERN and Pe are thought to be part of a neural error processing system that helps optimize actions and learning, as reflected in post-error adjustments of behavior. Such post-error adjustments include post-error slowing (PES) and post-error improvement in accuracy (PIA) (Danielmeier & Ullsperger, 2011). PES is thought to represent compensatory behavior where the RT is increased in the trial following an error in order to improve chances for accurate responding (Ullsperger et al., 2014). Another possibility is that it reflects the participant halting due to an automatic orienting response caused by an error or other surprising event (Danielmeier & Ullsperger, 2011; Wessel, 2018), or due to a greater perceived conflict in the preceding trial, as in post-conflict slowing (Bissett & Logan, 2011). To what degree PES reflects a conscious and strategic adaptation, or an automatic, unconscious reaction is debated (Danielmeier & Ullsperger, 2011), as is to what degree it is adaptive (Wessel, 2018). The developmental trajectory of PES is also unclear. Smulders et al. (2016) concluded that earlier findings were conflicting, with both increases, decreases and no changes in PES during childhood and adolescence being reported. As for PIA, Wessel (2018) postulates that it only occurs for trials with large inter-trial intervals of a second or more, with accuracy instead decreasing on post-error trials if the interval is short. Developmental studies on PIA are lacking. There is, however, evidence from adults to suggest that the degree of PIA is influenced by ERN and Pe amplitude, with greater amplitudes associated with increased PIA (Carp & Compton, 2009; Falkenstein et al., 2000). Similarly, some studies have found a stronger ERN to be associated with greater PES (Debener et al., 2005a; Gehring et al., 1993; Wessel & Ullsperger, 2011).

The neural underpinnings of electrophysiological error-processing components have been investigated using a range of different methods. A systematic comparison by Agam et al. (2011) of magnetoencephalography and high density electroencephalography (EEG) studies concluded that the cingulate cortex was the most likely source of the ERN, with the mean source locus between studies being in the dorsal anterior cingulate cortex (ACC). The exact locus varied substantially between studies, with several identifying the posterior cingulate cortex (PCC) as a source. Similarly, the ACC has been identified as the source in adolescents (Buzzell et al., 2017; Ladouceur et al., 2007; Santesso & Segalowitz, 2008). Single-unit recording in the ACC has corroborated it as a source of error or conflict processing in monkeys (Ito et al., 2003; Niki & Watanabe, 1979), and the same has been observed in humans undergoing cingulotomy (Davis et al., 2005; Sheth et al., 2012). Though the ACC seems to be a main source of the recorded ERN, the underlying latent mechanism seems to be dependent on a larger network, including limbic, subcortical, motor and prefrontal regions (Bush et al., 2000; Gehring et al., 2012; Huster et al., 2011a). Indeed, patients with ACC lesions have been shown to not produce an ERN, but nonetheless show error awareness (Stemmer et al., 2004). The evidence regarding the source of the Pe is less clear than for the ERN, but again the cingulate cortex seems to be the most likely candidate, albeit possibly with a more rostral source than the ERN (Herrmann et al., 2004; Veen & Carter, 2002). The surface area of the cingulate cortex, and most other cortical regions, shows the greatest age-related increase before approximately age 10 (Amlien et al., 2014), but relatively little change through adolescence (Fjell et al., 2018). Based on this, differences in cingulate cortex surface area in adolescence may not primarily reflect maturational differences, but rather more stable individual differences. In contrast, cortical thickness shows continued marked decrease with increasing age through both childhood and adolescence (Amlien et al., 2014; Tamnes et al., 2017; Vijayakumar et al., 2016; Wierenga et al., 2014).

In the present study, we investigated age-related differences in ERN and Pe amplitude during childhood and adolescence, and the associations between these brain functional indices of error processing and task performance, behavioral post-error adjustments and regional thickness and area of the cingulate cortex. Specifically, we analyzed EEG recorded during a Flanker task and structural magnetic resonance imaging (MRI) data from a separate session from participants 8-19 years old. Based on earlier studies (Grammer et al., 2014; Tamnes et al., 2013), we expected ERN amplitude to be greater for the older adolescents, while we expected Pe amplitude to show no relationship with age. Further, we expected ERN and Pe amplitude to be associated with task performance and with post-error adjustments, specifically that a stronger ERN would be associated with both greater PES and PIA and a stronger Pe be associated with a greater PIA, as observed previously in adults (Carp & Compton, 2009; Debener et al., 2005a; Gehring et al., 1993; Wessel & Ullsperger, 2011). Finally, we tentatively hypothesized that ERN and Pe amplitude would be negatively associated with cingulate cortex thickness and positively associated with cingulate cortex area, given the different developmental trajectories of these structural measures across adolescence. This is based on the assumption that a stronger ERN or Pe is generally indicative of more efficient processing and that a smaller cortical thickness may be indicative of greater relative maturity in the age group studied (Amlien et al., 2014), while greater cortical surface area is generally thought to be beneficial when looking at individual cognitive differences, including during development (Walhovd et al., 2016).

## 2. Materials and Methods

### 2.1 Participants

Participants between 8 and 19 years of age were recruited to the research project *Neurocognitive Development* (Østby et al., 2009; Tamnes et al., 2010) through newspaper advertisements, and local schools and workplaces. The study was approved by the Norwegian Regional Committee for Medical and Health Research Ethics. Written informed consent was provided by all participants over 12 years, and from a parent or guardian of participants younger than 18 years. Oral informed assent was obtained from participants younger than 12 years. Participants aged 16 years or older and a parent completed standardized health interviews regarding each participant. Exclusion criteria included premature birth (<37 weeks of gestation), a history of brain injury or disease, ongoing treatment for a mental disorder, use of psychoactive drugs, and MRI contraindications. Participants were required to be right-handed, fluent Norwegian speakers, and have normal or corrected-to-normal hearing and vision. A total of 113 children and adolescents fulfilled these criteria and were deemed free of significant brain injuries or conditions by a neuroradiologist. Seven participants were excluded due to task-performance criteria, as described in the next section. Further, eight more participants were excluded from the EEG analyses, and another seven participants from the MRI analyses due to poor data quality, as described in separate sections below. Age, sex and estimated IQ for each subsample are reported in **Table 1**. The four-subtest form of the Wechsler Abbreviated Scale of Intelligence was used to estimate IQ (Wechsler, 1999).

**Table 1.** Demographics for the different subsamples used for analyses using only behavioral data, both behavioral and EEG data, and both EEG and MRI data, respectively.

| Sample descriptives |  |  |  |
| --- | --- | --- | --- |
|  | Valid behavior | Valid behavior and EEG | Valid behavior, EEG and MRI |
| N | 106 | 98 | 92 |
| Age mean (SD) | 14.0 (3.4) | 14.1 (3.4) | 14.3 (3.3) |
| Age range | 8.3-19.7 | 8.3-19.7 | 8.4-19.7 |
| Sex | 54 f / 52 m | 48 f / 50 m | 46 f / 46 m |
| IQ mean (SD) | 109.2 (10.1) | 109.1 (10.0) | 109.2 (9.8) |

**Table 2.** Partial correlation for various ERP measures with age, controlling for RMS, and behavior, controlling for RMS and age. Significant correlations are marked with a star. Includes the area around the peak and delta waves used in the main text, as well as peaks of error trials with peaks of correct trials regressed out. Also included are the same analyses for the CRN, including the CRN peak with the ERN regressed out.

|  | ERN |  |  | Pe |  |  | CRN |  |
| --- | --- | --- | --- | --- | --- | --- | --- | --- |
|  | Peak | Delta | Residual | Peak | Delta | Residual | Peak | Residual |
| Age | -.21* | -.13 | -.19 | .05 | -.03 | -.08 | -.14 | -.10 |
| Accuracy | -.26* | -.23* | -.27* | .16 | .30* | .06 | -.02 | .04 |
| Reaction Time | .31* | .22* | .29* | .05 | -.16 | .06 | .12 | .06 |
| PES | -.23* | -.24* | -.25* | .06 | .19 | .06 | .03 | .05 |
| PIA | -.27* | -.28* | -.29* | .21* | .29* | .21* | .08 | .11 |

### 2.2 Stimuli and Task

Stimuli were presented on a 19-inch computer screen with a viewing distance of approximately 80 cm. Administrating and recording of the task was done with E-prime software using a Psychology Software Tools Serial Response Box. EEG was recorded during a modified and speeded version of the Eriksen arrow Flanker task (**Figure 1**), as previously described elsewhere (Tamnes et al., 2012). Briefly, stimuli were 2.5° vertical stacks of five 1 cm long white arrows, presented on a black background. Each trial started with a fixation cross presented for a random interval between 1200 and 1800 ms. Then, four arrows were presented for 80 ms before the target arrow was presented in the middle, together with the flanker arrows, for 60ms. This was done to make the task more difficult by priming for the prepotent response. Finally, a black screen was presented for up to 1440 ms. A total of 416 trials were presented, half of which were congruent, with the target arrow pointing in the same direction as the flankers, and the other half incongruent, with the target and flanker arrows pointing in opposing directions. Left and right pointing target arrows were equally frequent within each condition. Trials were presented in semi-random order, with incongruent trials never appearing more than three times in a row. Participants were asked to respond by pressing one of two buttons depending on the direction of the target arrow, using the index finger of each hand. They were also asked to emphasize both speed and accuracy when responding. An individual response time threshold was set for each participant based on their average reaction time on the first 20 trials. If participants responded slower than this threshold on three subsequent trials, they were asked to respond faster through a 1 second text prompt. This was done to increase error rate. There was a short break half-way through the task. Before the task, participants completed two practice blocks of 12 trials each. In the first of these, both the flankers and the target arrow were presented for slightly longer (150 and 90 ms, respectively), whereas in the second block, the times were the same as in the task.

**1.**
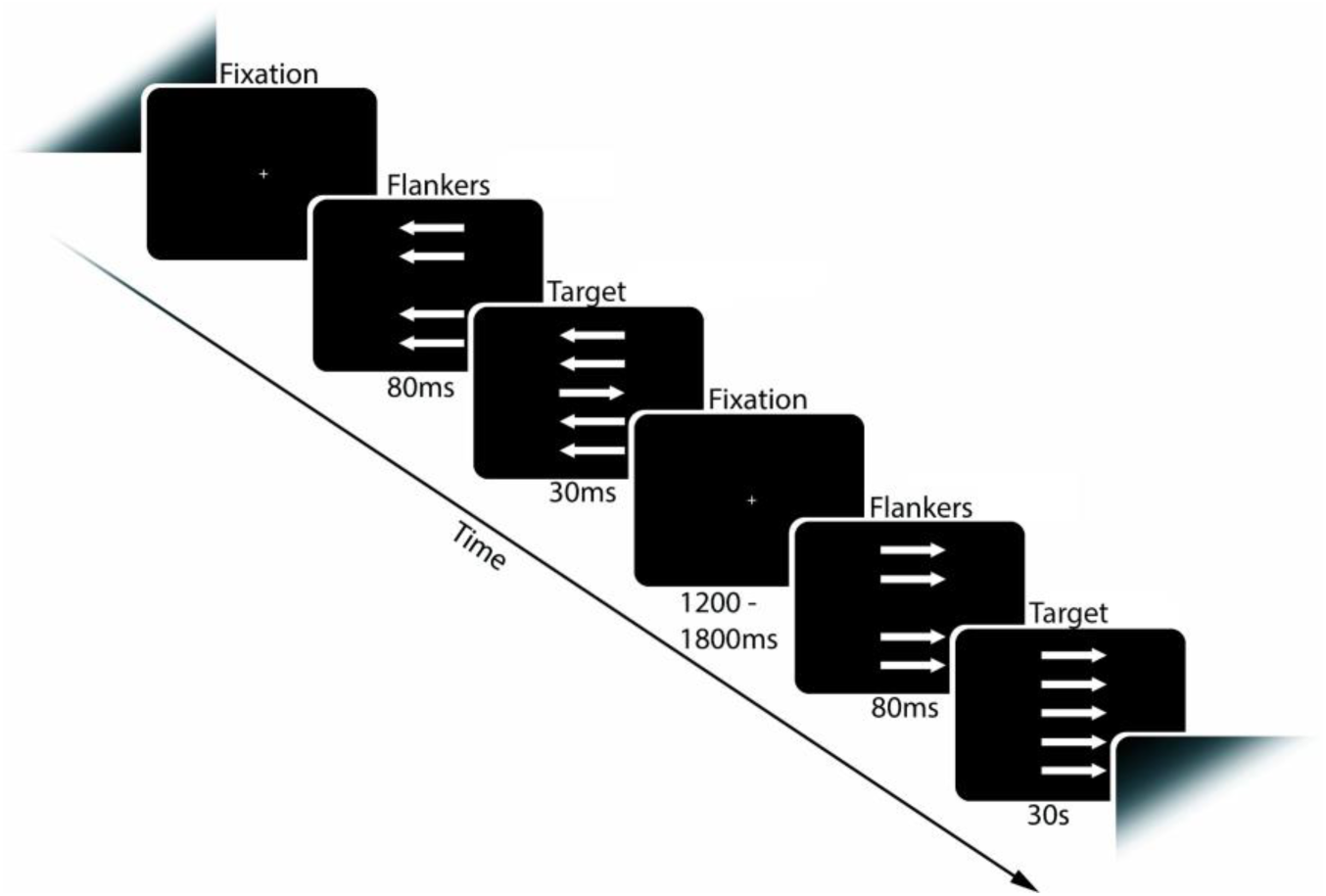
Flanker task. Schematic illustration of the task employed.

We used the following three behavioral inclusion criteria: First, participants were required to have ≥60% accuracy on congruent trials. Second, participants needed to show a significant congruency effect, with reaction times on correct incongruent trials significantly slower than on correct congruent trials as determined by a paired samples t-test (p <.05). These criteria were selected to exclude those with suboptimal motivation; not paying attention to the task or responding at random. Third, a minimum of six incongruent errors was required, which was estimated by Pontifex et al. (2010) to be the minimum number of required trials to get an internally consistent ERN and Pe in both children and adults. Three participants were excluded due to low accuracy, two more due to a lack of a congruency effect, and two more from having too few errors, yielding a sample of 106 for the behavioral analyses.

Median reaction time was calculated separately for correct and incorrect congruent and incongruent trials for each participant. Mean accuracy was calculated separately for congruent and incongruent trials for each participant. PES was calculated for each participant by subtracting the median reaction time of correct trials following correct trials from the median reaction time of correct trials following error trials. Similarly, PIA was calculated by subtracting the proportion of correct responses on trials following correct trials from the proportion of correct responses on trials following error trials. For follow-up analyses, we additionally calculated PES and PIA only for trials following incongruent trials to exclude the possible influence of post-conflict adjustments.

### 2.3 EEG Acquisition

Participants performed the task in an electrically shielded room while seated in a comfortable high-back chair. The electrophysiological recordings were done using 128 EEG channels with an electrode placement based on the 10% system (EasyCap Montage No. 15, http://www.easycap.de/). The sampling rate during recording was set to 1000 Hz. The electrodes used were EasyCap active ring electrodes (Ag/AgCl) with impedance conversion circuits integrated into the electrode housing that allows high quality recordings even with high impedance values, thus reducing preparation time and noise. The signals were amplified via a Neuroscan SynAmps2 system and filtered online with a 40 Hz low-pass and a 0.15 Hz high-pass analog filter prior to digitization and saving of the continuous data set. During recording, all electrodes were referenced to an electrode placed on the left mastoid. Eye blinks were recorded with one electrode above and one electrode below the left eye, and a ground electrode was placed anteriorly on the midline.

### 2.4 EEG Processing

Data pre-processing was done using Matlab and EEGLab. Bad channels, with insufficient or corrupt data, were identified using the clean_rawdata plugin (http://sccn.ucsd.edu/wiki/Plugin_list_process). Channels were rejected if they at any point during the recording flatlined for more than 5 seconds, if the channel correlated less than 0.85 with a reconstruction of this channel based on the surrounding electrodes, or if the channel had line noise relative to its signal that was four standard deviations or greater than the total channel population. Bad channels were interpolated from surrounding electrodes using EEGLab’s spherical interpolation, based on the method developed by Perrin et al. (1989). The data were segmented into response-locked 2000-ms epochs, starting 1000 ms before responses. Epochs were baseline-corrected relative to the time window 600-400 ms before responses, in order to avoid interference from the stimulus-elicited P300. Additionally, stimulus-locked epochs were extracted in 1600 ms epochs, starting 200 ms before presentation of the target arrow, and baseline corrected to the time window 200-100 ms before the target stimulus. Eye blink activity was identified and removed using EEGLab’s default independent component analysis (ICA) method. Activity tied to the stimulus was reduced from the response-locked averages using ADJAR-level 1 (Woldorff, 1993). In this procedure, for each participant, the average stimulus-locked activity is shifted over and subtracted from the average response-locked activity, based on and weighted by the reaction-time distribution of that participant.

Only incongruent error trials were used for peak extraction. This was done to control for between-subject variability in the proportion of incongruent and congruent error trials, keeping the relative interference from congruence effects consistent. All such epochs remaining after preprocessing were used. Peaks of the ERPs were identified individually for each participant. Peak latency of the ERN was defined for the combined channels FCz, Fz, FFC1h and FFC2h as the timing of the negative peak in the 0-100 ms time window following responses. Latency of the Pe was defined for the combined channels CPz, Pz, CPP1h and CPP2h as the timing of the positive peak in the 140-400 ms time window after responses. Channels and time windows were selected based on the known topography and timing of the ERPs of interest. Peak amplitudes for the ERN and the Pe were extracted as the average area under the curve of the 40 ms time window surrounding each peak, using the same channels that were used for determining peak latency. The terms peak and peak amplitude are used for these measures throughout the manuscript. Though not extracted from a single data point, which is the historically more common approach, the area around the peak represents a theoretically similar construct. For all analyses using our ERP measures, follow-up analyses were also performed using difference waves (ΔERN and ΔPe). These were calculated by subtracting the response-locked incongruent correct trials from the incongruent error trials. For each participant, an equal number of correct and error trials were used to generate ΔERN and ΔPe, with the number determined by whichever of the two conditions had the fewest trials, with trials from the more common condition selected randomly. ΔERN and ΔPe peaks were extracted for the same combined channels and time windows as the regular ERPs. These follow-up analyses were performed in order to have potentially purer measures of error processing, with variance shared between correct and error trials filtered out, as well as to make the results directly comparable to previous developmental studies reporting on these measures. As a supplementary analysis we also looked at the ERN and Pe with activity from the correct trials regressed out in relation to age and behavior. We examined the CRN in the same fashion. These analyses are described in the supplementary materials.

### 2.5 MRI Acquisition

A 1.5 Tesla Siemens Avanto scanner (Siemens Medical Solutions) with a 12-channel head coil was used to acquire MRI data. For the morphometric analyses we used a 3D T1-weighted MPRAGE pulse sequence with the following parameters: TR/TE/TI/FA = 2400 ms/3.61 ms/1000 ms/8°, matrix 192 × 192, field of view = 240, 160 sagittal slices, voxel size 1.25 × 1.25 × 1.20 mm. Duration of the sequence was 7 min 42 s. A minimum of two repeated T1-weighthed sequences were acquired. All images were screened immediately after data acquisition and rescanning was performed if needed and possible. The protocol also included a 176-slice sagittal 3D T2-weighted turbo spin-echo sequence (TR/TE = 3390/388 ms) and a 25-slice coronal FLAIR sequence (TR/TE = 7000–9000/109 ms) to aid the radiological examination.

### 2.6 MRI Processing

For each participant, the T1-weighthed sequence with best quality as determined by visual inspection of the raw data was chosen for analysis. Whole-brain volumetric segmentation and cortical reconstruction was performed with FreeSurfer 5.3 (http://surfer.nmr.mgh.harvard.edu/). The details of the procedures are described elsewhere (Dale et al., 1999; Fischl, 2012; Fischl et al., 2002; Fischl et al., 1999). Cortical surface area (white matter surface) maps were computed by calculating the area of every triangle in the tessellation. The triangular area at each location in native space was compared with the area of the analogous location in registered space to give an estimate of expansion or contraction continuously along the surface (“local arealization”) (Fischl et al., 1999). Cortical thickness maps for each participant were obtained by calculating the distance between the cortical gray matter and white matter surface at each vertex (Fischl & Dale, 2000). All processed scans were visually inspected in detail for movement and other artifacts. The cortical surface was parcellated into 33 different gyral-based regions in each hemisphere (Desikan et al., 2006), of which three regions of the cingulate cortex were selected as regions of interest (ROI): rostral ACC (rACC), caudal ACC (cACC) and PCC (**2**). Surface area and mean cortical thickness of these ROIs in each hemisphere were used for further analyses.

### 2.7 Statistical Analysis

Descriptive statistics and Pearson’s correlation analyses were used to characterize the sample and task performance, and to test how task performance and post-error adjustments were associated with age. In all analyses that included ERP measures, only data from incongruent trials were used. To correct for potential noise differences in EEG data between participants caused by a variable number of error trials being used, the root-mean square (RMS) was calculated for the baseline of the averaged incongruent error trial for each participant. In all analyses involving ERPs, this noise estimate was included as a covariate. Partial correlations, controlling for RMS, were used to assess the relationships between ERP amplitudes and age, while partial correlations, controlling for age and RMS, were used to assess the relationships between ERP amplitudes and task performance and post-error adjustments. In order to explore interactions between age and ERP measures on accuracy, reaction time and post-error effects, multiple linear regression analyses, controlling for age and RMS, were performed. Partial correlations, controlling for age were used to assess the correlation between PES and PIA.

Pearson’s correlation analyses were used to explore the associations between age and cortical thickness and surface area in the selected cingulate ROIs. Partial correlations, controlling for age, sex and RMS, were used to assess the relationship between ERP amplitudes and regional cingulate cortex thickness and area. For the analyses involving cortical thickness or surface area, we controlled for multiple comparisons using a Bonferroni correction procedure adjusted for correlated variables (http://www.quantitativeskills.com/sisa/calculations/bonfer.htm) (Perneger, 1998; Sankoh et al., 1997). As a follow-up, surface-based whole-brain cortical analyses were performed vertex-wise (point-by-point) using general linear models, as implemented in FreeSurfer 6.0. Main effects of ERN and Pe peak amplitude on cortical structure were tested, while controlling for the effects of sex, age and RMS. Separate analyses were performed for cortical surface area and thickness maps. The data were tested against an empirical null distribution of maximum cluster size across 10,000 iterations using Z Monte Carlo simulations as implemented in FreeSurfer (Hagler et al., 2006; Hayasaka & Nichols, 2003) synthesized with a cluster-forming threshold of P < .001, yielding clusters corrected for multiple comparisons across the surfaces. Cluster-wise corrected P < .01 was regarded significant.

## 3. Results

### 3.1 Task Performance

Included participants responded accurately on 94.8% (SD = 6.8%) of congruent trials and 73.2% (SD = 15.0%) of incongruent trials. Mean reaction times in the congruent condition were 391 ms (SD = 87) for correct responses and 326 ms (SD = 111) for error responses, while in the incongruent condition they were 485 ms (SD = 76) for correct responses and 321 ms (SD = 68) for error responses. Accuracy for congruent trials correlated positively with age (r= .53, p < .001), while accuracy for incongruent trials did not (r = .04, p = .676). Reaction times were negatively correlated with age for both incongruent correct trials (r = −.68, p < .001), incongruent errors trials (r = −.55, p < .001), and congruent correct trials (r = −.67, p < .001), but not for congruent incorrect trials (r = −.13, p = .214).

### 3.2 Age-related Differences in ERN and Pe

The ERN was identified as a sharp, frontal negative deflection peaking on average 32 ms post response, while Pe was identified as a broader, centro-parietal positive deflection peaking at 196 ms post response (Figure **3**). Controlling for RMS, amplitude of the ERN correlated negatively with age (r = −.21, p = .043), indicating a slightly stronger negative amplitude with older age. There was no correlation between Pe amplitude and age (r = −.13, p = .203). For the difference waves between incongruent error and correct responses, there were no correlations between age and ΔERN amplitude (r = −.05, p = .629) or age and ΔPe amplitude (r = −.03, p = .773).

**Figure 2.**
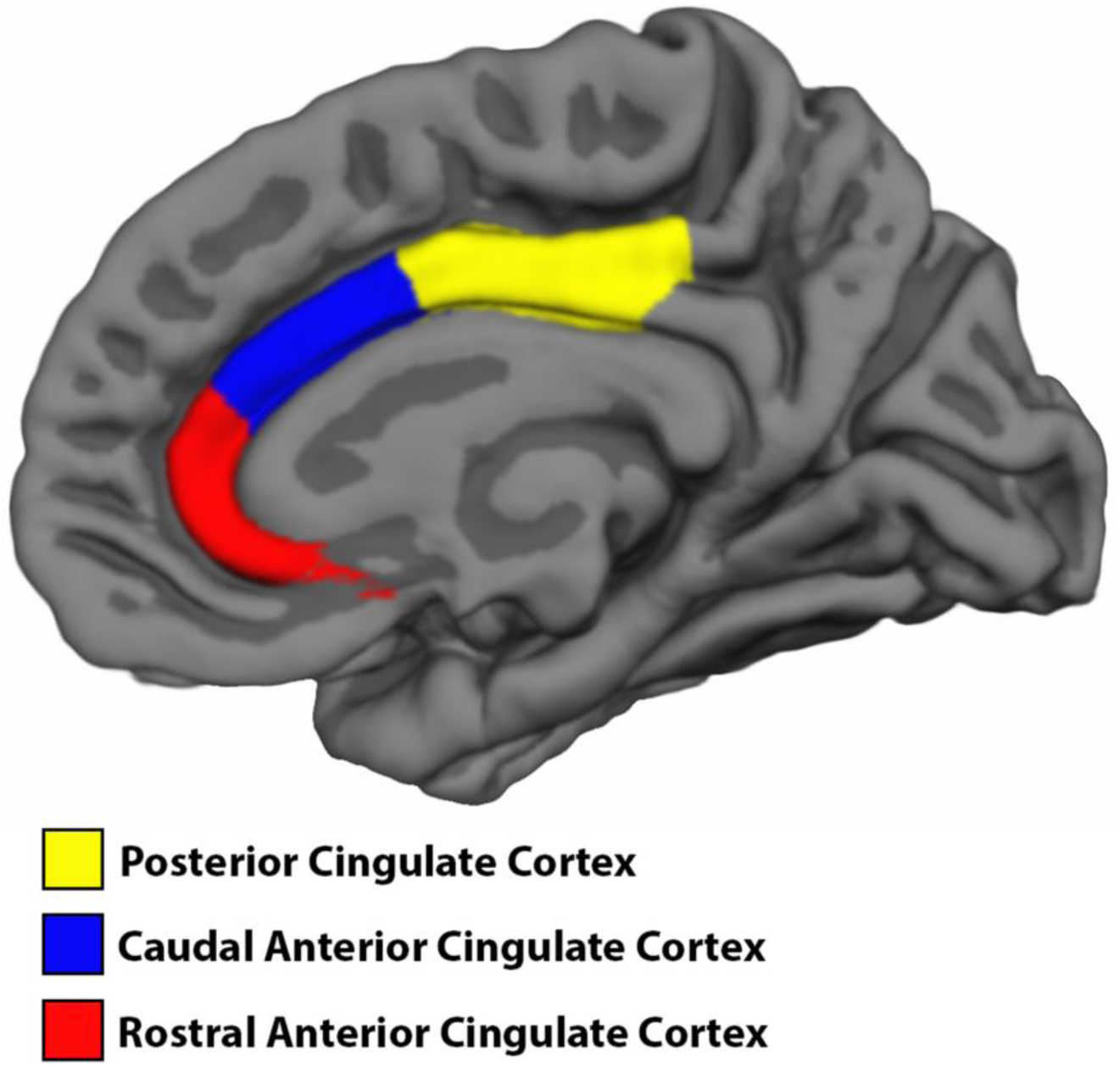
Extracted Cingulate Cortex Regions of Interest. Color coded Regions of Interest of the cingulate cortex used in cortical analyses. Includes Posterior Cingulate Cortex (yellow), Caudal Anterior Cingulate Cortex (blue) and Rostral Anterior Cingulate Cortex (red).

**Figure 3.**
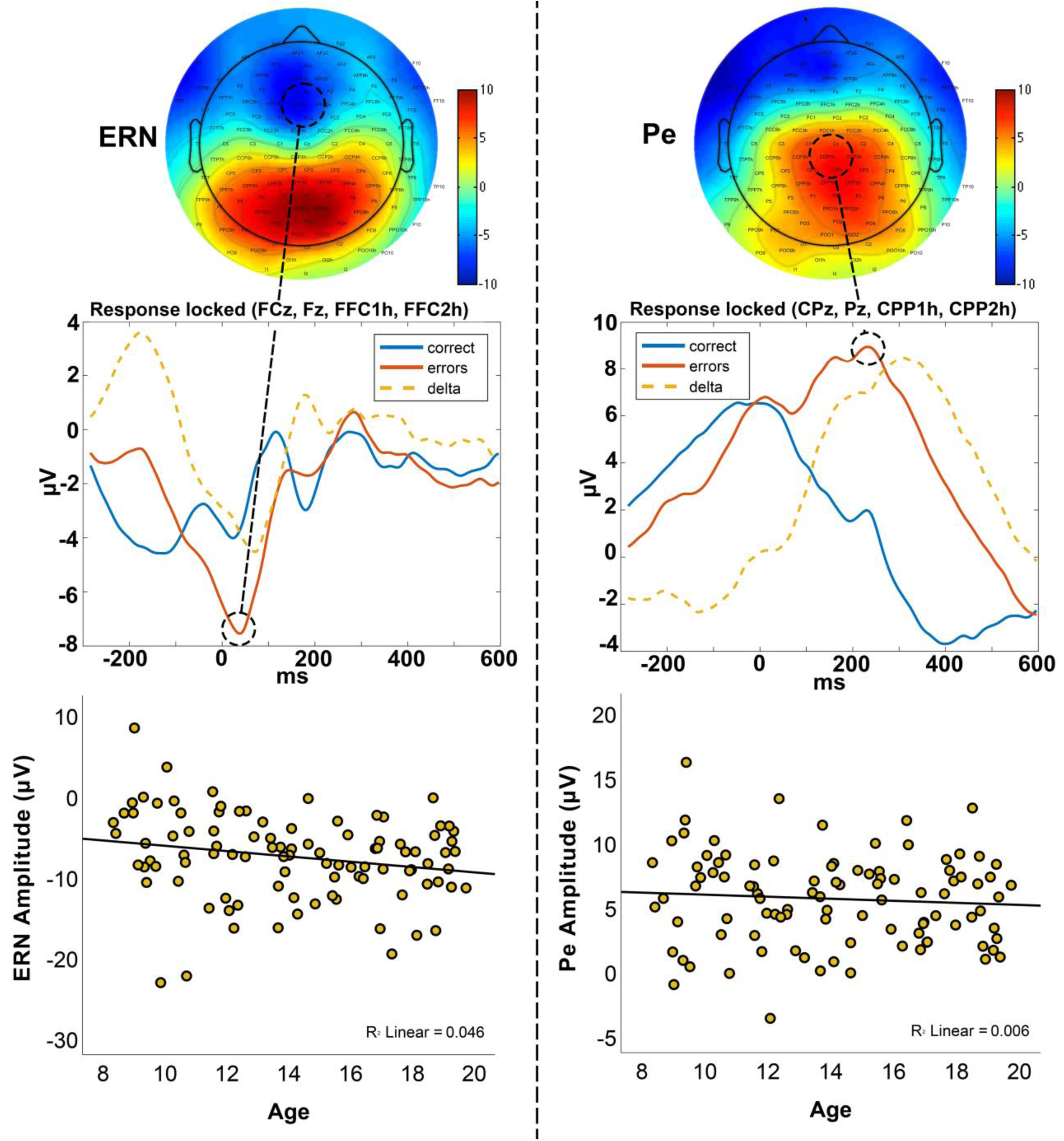
Electrophysiological markers of error processing. Scalp topographies (top), grand averaged response-locked time courses (middle) and scatter plots showing associations with age (bottom) for the ERN (left) and Pe (right). Time courses are split between incongruent correct trials (blue), incongruent error trials (red) and the difference, or delta, between the two (orange). Data in the scatter plots are from incongruent trials and µV is here the mean voltage of a 40 ms time window surrounding each peak.

**Figure 4.**
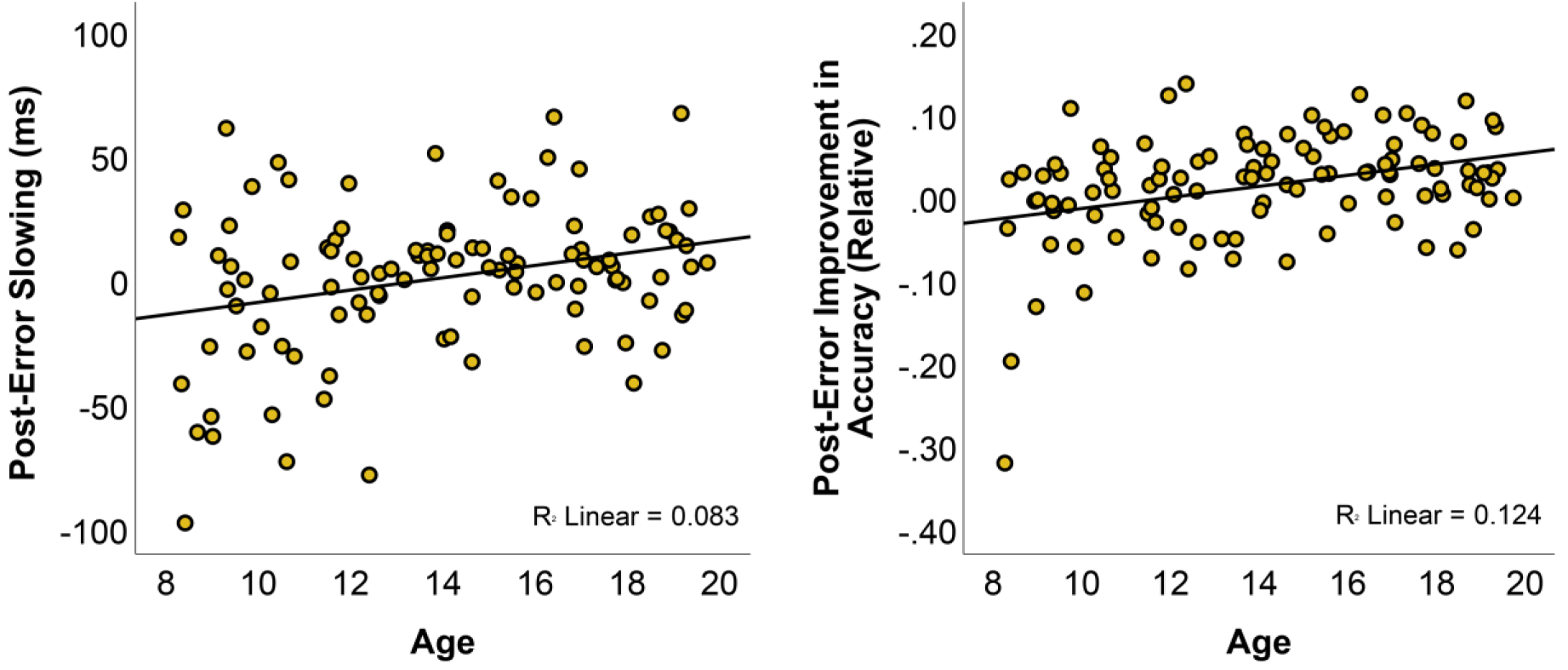
Associations between post-error adjustments and age. Scatter plots showing the relationships between age and PES (left) and PIA (right).

### 3.3 Associations between ERN and Pe and Task Performance

Associations between ERPs and task performance were tested with partial correlations, controlling for age and RMS. There were significant negative correlations between accuracy and both ERN amplitude (r = −.26, p = .012) and ΔERN amplitude (r = −.23, p = .024), indicating that a stronger ERN was associated with better behavioral performance. There were also significant correlations between reaction time on correct incongruent trials and both ERN amplitude (r = .31, p = .002) and ΔERN amplitude (r = .22, p = .030). For Pe, there was no significant correlation between accuracy and Pe amplitude (r = .16, p = .118), but accuracy was significantly correlated with ΔPe amplitude (r = .30, p = .003), indicating that specifically stronger Pe was associated with higher accuracy. Reaction time on correct incongruent trials was not associated with Pe amplitude (r = 05, p = .624) or ΔPe amplitude (r = −.16, p = .120). Multiple regression analyses revealed several significant interactions between age and ERP peaks for the behavioral criterion variables. There were significant interactions between ERN amplitude and age on PES (Beta = 1.06, t(97) = 2.51, p = .014) and PIA (Beta = 1.10, t(97) = 2.71, p = .008). In both cases this indicated a stronger association between ERN amplitude and post-error adjustment with higher age. No significant interactions were found between ERN and age on accuracy (Beta = −.61, t(97) = −1.46, p = .147) or reaction time (Beta = −.49, t(97) = −1.67, p = .099). There were significant interactions between Pe amplitude and age on accuracy (Beta = −.93, t(97) = −2.41, p = .018), which indicated a decreased association between Pe amplitude and accuracy with age. For Pe, no other significant age interactions were found, either on reaction time (Beta = −.51, t(97) = −1.77, p = .080), PES (Beta = −.31, t(97) = −.76, p = .451) or PIA (Beta = −.51, t(97) = −1.32, p = .190).

### 3.4 Post-error Adjustments

There were significant positive correlations between age and both PES (r = .29, p = .003) and PIA (r = .35, p < .001) (**4**), indicating that older participants were relatively slower and more precise on trials following errors compared with younger participants. There was also a correlation between PES and PIA, even when controlling for age (r = .24, p = .015). Partial correlations, controlling for age and RMS, were performed to test for age-independent associations between ERP amplitudes and post-error adjustments. Amplitude of the ERN showed a negative correlation with PES (r = −.24, p = .017), but not with PIA (r = −.17, p = .101), though ΔERN amplitude showed negative correlations with both PES (r = −.21, p = .037) and PIA (r = −.27, p = .009). Pe amplitude was not associated with either PES (r = .05, p = .660) or PIA (r = .108, p = .294). The same was true for ΔPe amplitude (PES: r = .18, p = .073; PIA: r = .18, p = .086).

In follow-up analyses, we repeated the above analyses with PES and PIA calculated only for trials following incongruent trials to exclude the possible influence of post-conflict adjustments. Here, there was still a positive correlation between age and PES (r = .20 p = .040), but not with PIA (r = .16 p = .099), and PES and PIA no longer showed an age-independent association (r = .17 p = .080). For associations with ERPs, there was still a significant association between ERN and PES (r = −.218 p = .033).

### 3.5 Associations between ERN and Pe and Cingulate Cortex Thickness and Area

The average correlation between the six included structural MRI ROIs (rACC, cACC and PCC in each hemisphere) was r = 0.34 for cortical thickness and r = 0.42 for surface area. Using a Bonferroni correction procedure adjusted for correlated variables this yielded corrected significance levels at .015 for thickness and .018 for area. There were negative correlations between age and cortical thickness for all regions; rACC (LH: r = −.53, p < .001; RH: r = −.34, p < .001), cACC (LH: r = −.32, p < .001; RH: r = −.37, p < .001) and PCC (LH: r = −.49, p < .001; RH: r = −.51, p < .001). No significant correlations were found between age and cortical surface area for any of the ROIs after controlling for multiple comparisons. Partial correlations were performed to test the associations between ERP amplitudes (ERN, Pe, ΔERN, ΔPe) and the thickness and surface area of the three regions of the cingulate cortex for each hemisphere, controlling for sex, RMS and age. There were no corrected significant associations. Follow-up analyses using whole-brain vertex-wise GLMs also yielded no significant clusters for either ERN or Pe amplitude.

## 4. Discussion

In this study, we found an association between ERN amplitude and age, with older adolescents having a more negative ERN than younger individuals. In contrast, Pe amplitude was not associated with age. This is generally in line with earlier research (Davies et al., 2004b; Hogan et al., 2005; Ladouceur et al., 2007; Wiersema et al., 2007), and supports the notion that the Pe reflects a process that largely matures in childhood (Wiersema et al., 2007), while the ERN reflects a process that continues to refine in adolescence (Tamnes et al., 2013). We also found that a stronger ERN was associated with higher task accuracy and faster reaction times, and that a stronger Pe was associated with higher accuracy. Moreover, both PES and PIA increased with age, and a stronger ERN amplitude was associated with greater PES. We found no relationships between the ERPs and regional cingulate cortex thickness or surface area.

Behaviorally, older adolescents showed faster response times compared with younger participants. Task accuracy was positively associated with age for congruent, but not incongruent trials. The ERN was identified frontally on the scalp and had a sharp peak shortly after the target presentation. The Pe was identified at centro-parietal recording sites, and had a wider, less defined peak. Amplitude of the ERN was more negative for older participants, while Pe amplitude was not associated with age. What these associations tell us about cognitive development is dependent on what functions we attribute to the ERN and Pe. If we use the interpretation that the Pe indexes awareness and strategic adjustment (Nieuwenhuis et al., 2001; Ridderinkhof et al., 2009), while the ERN reflects more automatic processing, the differing trajectories would imply that while the ability to consciously detect and process conflict-inducing events such as errors reaches full maturity early, the ability to respond quickly and automatically to such events has a more protracted development. Interestingly, these age-associations differ from what has been observed for the related N200 and P300 ERPs, where the N200 has generally been shown to decrease in strength with age (Downes et al., 2017), and P300 to increase (van Dinteren et al., 2014), which is also what we have observed previously in the same sample as the present study (Overbye et al., 2018).

Both ERN and Pe amplitude were, independently of age, associated with task accuracy, with stronger amplitudes related to higher accuracy. This has previously been observed in adults for ERN (Westlye et al., 2008), and in children for both ERN and Pe (Torpey et al., 2012). Also, a more negative ERN was associated with faster reaction times in our sample, consistent with earlier research on children (Torpey et al., 2012). Together, these results fit with the interpretation that both the ERN and Pe reflect processes that are functionally relevant for optimizing actions, and ultimately learning.

When analyzing correct trials following both congruent trials and incongruent trials, both PES and PIA were positively associated with age, i.e. greater for older adolescents compared with younger participants. In other words, older adolescents responded to errors by slowing down and improving accuracy to a greater degree than did younger participants. A positive association between PES and age has been shown in one previous developmental study (Hogan et al., 2005), while others have shown the opposite trend (Taylor et al., 2018) or more complex patterns (Ladouceur et al., 2007). The function of PES is a debated subject, with hypotheses ranging from a tactical slowing in order to improve accuracy (Ullsperger et al., 2014), to a maladaptive orienting response to surprising, infrequent errors (Alexander & Brown, 2010; Danielmeier & Ullsperger, 2011). That PES is not always adaptive is supported by findings indicating that PES and PIA are not always related (Danielmeier & Ullsperger, 2011). In our results, however, PES and PIA correlate positively. A possible reconciliation of these findings is that PES is not a unitary construct, but might represent different constructs depending on the test paradigm. Wessel (2018) suggests that PES is a result of an immediate halting of an ongoing response following an error or surprising event, followed by a slower adaptation of behavior, meaning that PES will only be related to PIA if there is sufficient time following the error. In accordance with this, PIA is generally only seen at long response-stimulus intervals (RSI) (Danielmeier & Ullsperger, 2011; Wessel, 2018). Since we used long RSIs (between 1280 and 1880 ms) our results are generally in line with this hypothesis. Furthermore, a stronger ERN was associated with greater PES, similar to what has been observed in adults (Debener et al., 2005b; Gehring et al., 1993) and adolescents (Ladouceur et al., 2007). We additionally found ERN amplitude to have stronger associations with PES and PIA with higher age, which might indicate an increased ability of the maturing error processing system in guiding behavior.

Post-error adjustments are conceptually linked to post-conflict adjustments (Danielmeier & Ullsperger, 2011), which led us to perform follow-up analyses of PES and PIA where we controlled for the influence of conflict processing by only including trials following incongruent trials. We chose incongruent trials for this correction as incongruent errors were more frequent than congruent errors. Here, the association between age and PES after error in incongruent trials only was still significant, as was the association between ERN and PES, although these correlations were weakened. There was however no longer a significant association between age and PIA or between PIA and PES. This raises the question if some of the observed associations can be attributed to a post-conflict adjustments (Chang et al., 2014), rather than post-error adjustments, considering that participants varied in the proportion of congruent to incongruent errors. In general, conflict processing and error processing seem to involve similar processes, and the N200 and P300 ERPs following conflict mirror in many ways the ERN and Pe following error commission (Gruendler et al., 2011). Still, some argue that conflict and error processing are not entirely interchangeable, and are possibly mediated by somewhat different neural mechanisms (Chang et al., 2014). Despite these theoretical and methodological concerns, our results seem to indicate that the ability to adjust behavior in response to errors improves through adolescence, and that this ability is at least partly caused by slowing down responses. In a developmental fMRI study, Velanova et al. (2008) suggested that the late maturation of error regulation might be tied to immaturity of the dACC. They found that the activity of the dACC showed greater differentiation between errors and correct trials for adults compared to children and adolescents.

Several lines of evidence, mainly from studies on adults, as well as animal studies, show that the ERN and Pe are most likely generated in the cingulate cortex, with both the ACC and the PCC implicated (Agam et al., 2011; Herrmann et al., 2004; Tamnes et al., 2013). In the present study, we aimed to investigate the links between these electrophysiological markers of error processing and regional cortical thickness and area of the cingulate. Through adolescence, cortical thickness in general decreases substantially, while surface area remains relatively stable in comparison (Tamnes et al., 2017; Vijayakumar et al., 2016). Based on this, and studies indicating positive associations between cognitive performance and surface area (Curley et al., 2018; Fjell et al., 2012), we hypothesized that ERN and Pe amplitude would be negatively associated with cingulate cortex thickness, reflecting individual maturation differences even when statistically controlling for age, and positively associated with cingulate cortex area, reflecting stable individual differences. In adults, we have previously found associations between ERN amplitude and both white matter and intracortical microstructure in the left PCC (Grydeland et al., 2016; Westlye et al., 2008), and ERN amplitude has also been found to be positively associated with cortical volume in the left cACC (Araki et al., 2013). In youth, we recently found a positive relationship between P300 component strength and cortical surface area in a region in the left inferior temporal gyrus (Overbye et al., 2018), but no previous studies have tested the associations between ERN or Pe and brain structure. We did, however, not find any significant associations between either ERN or Pe and regional cingulate cortex thickness or surface area in the present study. Considering our modest sample size, and thus limited statistical power, future studies are needed to further investigate this. Moreover, although the cingulate cortex seems to be the main generator of both the ERN and Pe, with both the ACC and PCC being implicated, both ERPs depend on larger networks and the rapid communication between network nodes (Taylor et al., 2007). The ERN is also associated with activity in the basal ganglia and the presupplementary motor region (Huster et al., 2011b), while the Pe has been associated with parietal and prefrontal brain regions (Hester et al., 2005). Thus, it might be fruitful to explore the associations between ERN and Pe and structural measures of these regions, as well as with structural or functional connectivity between the implicated regions as inferred from diffusion MRI or functional MRI. Finally, future studies should, to a greater degree than what has been done up to this point, implement longitudinal designs to more directly investigate development and stability of the ERN and Pe. The cross-sectional design of our study makes it difficult to distinguish developmental differences from cohort effects. Another limitation is that we did not examine sex differences in error processing or its development. A recent study indicated that males have greater ERN amplitudes and possibly smaller PES compared to females (Fischer et al., 2016).

## 5. Conclusions

Our results suggest that the ERN amplitude grows stronger through adolescence, while the Pe amplitude remains relatively stable. Age-independent associations were found between the strength of these components and task performance. Moreover, the results suggested that both PES and PIA increase with age, although post-conflict adjustment appeared to account for the latter effect, and that a stronger ERN is associated with greater PES. We found no significant associations between the ERN or Pe and regional cingulate cortex thickness or surface area. Combined, our results provide a comprehensive cross-sectional description of the maturation of the error processing system, including central electrophysiological markers and behavioral post-error adjustments, across adolescence.

## Acknowledgments

This study was supported by the Research Council of Norway (to KBW and to CKT), and the University of Oslo (to KO).

## 6. Supplementary materials

### 6.1 Residualized scores and the CRN

Meyer et al. (2017) have suggested that when trying to isolate ERN activity, it might be beneficial to regress out the activity from correct trials rather than using a subtraction-based difference score. They argue that this approach creates purer measures of the activity specifically related to the processing of correct and incorrect responses, without introducing additional variance from the other condition. As a follow-up we reran our analysis comparing the ERP and Pe with age and behavior, using residualized scores. Table 2 shows the results of these new analyses as compared to the results using regular peaks and delta measures. The results for the residualized scores were overall similar to the results using regular peaks. While the residualized ERN, in contrast to the peak ERN, did not have a significant correlation with age, the correlation coefficients were highly similar. Otherwise, all previously significant correlations remain significant, while no new significant correlations appeared. No significant correlations were found for the CRN using either regular peaks or residualized scores. Seen together, these results indicate that the CRN has little predictive value regarding the variables of interest in this study and that the peak measures used are already a decent approximation of the latent process specific to error trials.

